# Reduced neutralization of SARS-CoV-2 variants by convalescent plasma and hyperimmune intravenous immunoglobulins for treatment of COVID-19

**DOI:** 10.1101/2021.03.19.436183

**Authors:** Juanjie Tang, Youri Lee, Supriya Ravichandran, Gabrielle Grubbs, Chang Huang, Charles Stauft, Tony Wang, Basil Golding, Hana Golding, Surender Khurana

**Author notes:** These authors contributed equally to this manuscript (Juanjie Tang, Youri Lee, Supriya Ravichandran, Gabrielle Grubbs). **Corresponding author:** Surender Khurana, Ph.D. Division of Viral Products, Center for Biologics Evaluation and Research (CBER), Food and Drug Administrationa (FDA), 10903 New Hampshire Avenue, Silver Spring, MD, 20993, USA, E. mail.

## Abstract

Hyperimmune immunoglobulin (hCoV-2IG) preparations generated from SARS-CoV-2 convalescent plasma (CP) are under evaluation in several clinical trials of hospitalized COVID-19 patients. Here we explored the antibody epitope repertoire, antibody binding and virus neutralizing capacity of six hCoV-2IG batches as well as nine convalescent plasma (CP) lots against SARS-CoV-2 and emerging variants of concern (VOC). The Gene-Fragment Phage display library spanning the SARS-CoV-2 spike demonstrated broad recognition of multiple antigenic sites spanning the entire spike including NTD, RBD, S1/S2 cleavage site, S2-fusion peptide and S2-heptad repeat regions. Antibody binding to the immunodominant epitopes was higher for hCoV-2IG than CP, with predominant binding to the fusion peptide. In the pseudovirus neutralization assay (PsVNA) and in the wild-type SARS-CoV-2 PRNT assay, hCoV-2IG lots showed higher titers against the WA-1 strain compared with CP. Neutralization of SARS-CoV-2 VOCs from around the globe were reduced to different levels by hCoV-2IG lots. The most significant loss of neutralizing activity was seen against the B.1.351 (9-fold) followed by P.1 (3.5-fold), with minimal loss of activity against the B.1.17 and B.1.429 (≤2-fold). Again, the CP showed more pronounced loss of cross-neutralization against the VOCs compared with hCoV-2IG. Significant reduction of hCoV-2IG binding was observed to the RBD-E484K followed by RBD-N501Y and minimal loss of binding to RBD-K417N compared with unmutated RBD. This study suggests that post-exposure treatment with hCoV-2IG is preferable to CP. In countries with co-circulating SARS-CoV-2 variants, identifying the infecting virus strain could inform optimal treatments, but would likely require administration of higher volumes or repeated infusions of hCOV-2IG or CP, in patients infected with the emerging SARS-CoV-2 variants.

## INTRODUCTION

An expedited access to treatment of COVID-19 patients with convalescent plasma was issued by FDA via Emergency Use Authorization on August 23, 2020. Additional studies, including randomized, controlled trials, have provided data to further inform the safety and efficacy of COVID-19 convalescent plasma. Based on assessment of these data, potential clinical benefit of transfusion of COVID-19 convalescent plasma in hospitalized patients with COVID-19 is associated with high neutralizing titer units administered early in the course of disease(*1, 2*).

Intravenous immunoglobulins (IVIG) are a more concentrated form of IgG preparations fractionated from large number of plasma units that are prescreened for the presence of high titer anti-spike antibodies and predetermined SARS-CoV-2 neutralization titers. Several hCoV-2IG lots are currently being evaluated in clinical trials. The effectiveness of hCoV-2 IG products may be hampered by evolving SARS-CoV-2 and the emergence of new variants with high transmissibility rates and mutations in the Receptor Binding Domain (RBD) which are less susceptible to antibodies from recovered COVID-19 patients. The main variants of concern (VOC) are the B.1.1.7 spreading from the UK, the B1.351 spreading in South Africa (SA), and the P.1 that appeared in northeast Brazil and found in Japan (JP). In the US, several variants were identified recently including California (CA) variant B.1.429 (*3–6*).

The phage display technique is suitable to display properly folded and conformationally active proteins, as it has been widely used for display of large functionally-active antibodies, enzymes, hormones, and viral and mammalian proteins. We have adapted this Genome Fragment Phage Display Library (GFPDL) technology for unbiased, comprehensive approach for multiple viral pathogens including SARS-CoV-2, Ebola virus, highly pathogenic avian influenza virus, respiratory syncytial virus and Zika virus, to define both the linear and conformational antibody epitope repertoire of post-vaccination/infection samples (*7–11*)

In the current study we probed the antibody epitope repertoires of 6 hCoV-2IG products using SARS-CoV-2-spike GFPDL. Surface Plasmon Resonance (SPR) was used to measure antibody binding to SARS-CoV-2 antigenic site peptides identified in the GFPDL analyses and to spike protein receptor binding domain (RBD) representing WA-1 as well RBD mutants engineered to express key amino acid mutations of the VOCs. Neutralization capacity of the hCoV-2IG lots against the SARS-CoV-2 WA-1 strain and several VOC (CA, UK, JP, SA) was measured in pseudovirion neutralization assay (PsVNA) as well as classical PRNT assay. For comparison with hCoV-2IG, we evaluated nine convalescent plasma from recovered COVID-19 patients and 16 IVIG preparations that were manufactured with pre-pandemic plasma units prior to August 2019.

## RESULTS

### SARS-CoV-2 spike antibody epitope repertoires of six hCoV-2IG batches

The spike protein is the antigen of choice for development of vaccines and therapeutics against SARS-CoV-2. To decipher the epitope-specificity of the SARS-CoV-2 spike-specific antibodies in an unbiased manner, we subjected the six hCoV-2IG lots to antibody epitope profiling with a highly diverse SARS-CoV-2 spike GFPDL with >10^7.1^ unique phage clones displaying epitopes of 18-500 amino acid residues across the SARS-CoV-2 spike. During GFPDL characterization, GFPDL based epitope mapping of monoclonal antibodies (MAbs) targeting SARS-CoV-2 spike or RBD identified the expected linear or conformation-dependent epitopes recognized by these MAbs. Recently, we showed that SARS-CoV-2 spike GFPDL can recognize both linear, conformational and neutralizing epitopes in the post-vaccination sera of rabbits (*11*) and post-SARS-CoV-2 infection sera in the adults and elderly (*10, 12*).

Six hCoV-2IG lots were used for SARS-CoV-2 GFPDL based epitope mapping as previously described (*7–11, 13*). Similar numbers of phages were bound by IgG of these hCoV-2IG batches (3.4 x 10^4^ – 2.1 x 10^5^) (Fig. 1A). The bound phages demonstrated a diverse epitope repertoire spanning the entire SARS-CoV-2 spike protein including N-terminal domain (NTD) and RBD in S1, and the fusion peptide (FP), β-rich connector domain (CD), heptad repeat 1 (HR1) and 2 (HR2) in S2 (Fig. 1B and Fig. S1). Peptides representing the key immunodominant antigenic sites identified by GFPDL analysis were chemically synthesized and used to evaluate binding of each of the six hCoV-2IG batches, 16 pre-pandemic 2019-IVIG lots and 9 COVID-19 convalescent plasma (Fig. 1C). As expected 2019-IVIG demonstrated minimal to no binding to the SARS-CoV-2 spike peptides. In aggregate, the convalescent plasma showed lower binding to epitopes spanning the entire spike in comparison with the hCoV-2IG (Fig. 1C). In agreement with GFPDL analysis, the hCoV-2IG demonstrated highest antibody binding to the spike peptide 790-834 that contains the fusion peptide sequence (residues 788-806), which is unchanged among the major VOCs ((Fig. S2 and Table S1). Most of the GFPDL-identified antigenic site sequences recognized by hCoV-2IGs are conserved in the spike protein of various SARS-CoV-2 VOC (Table S1).

**Figure 1.**
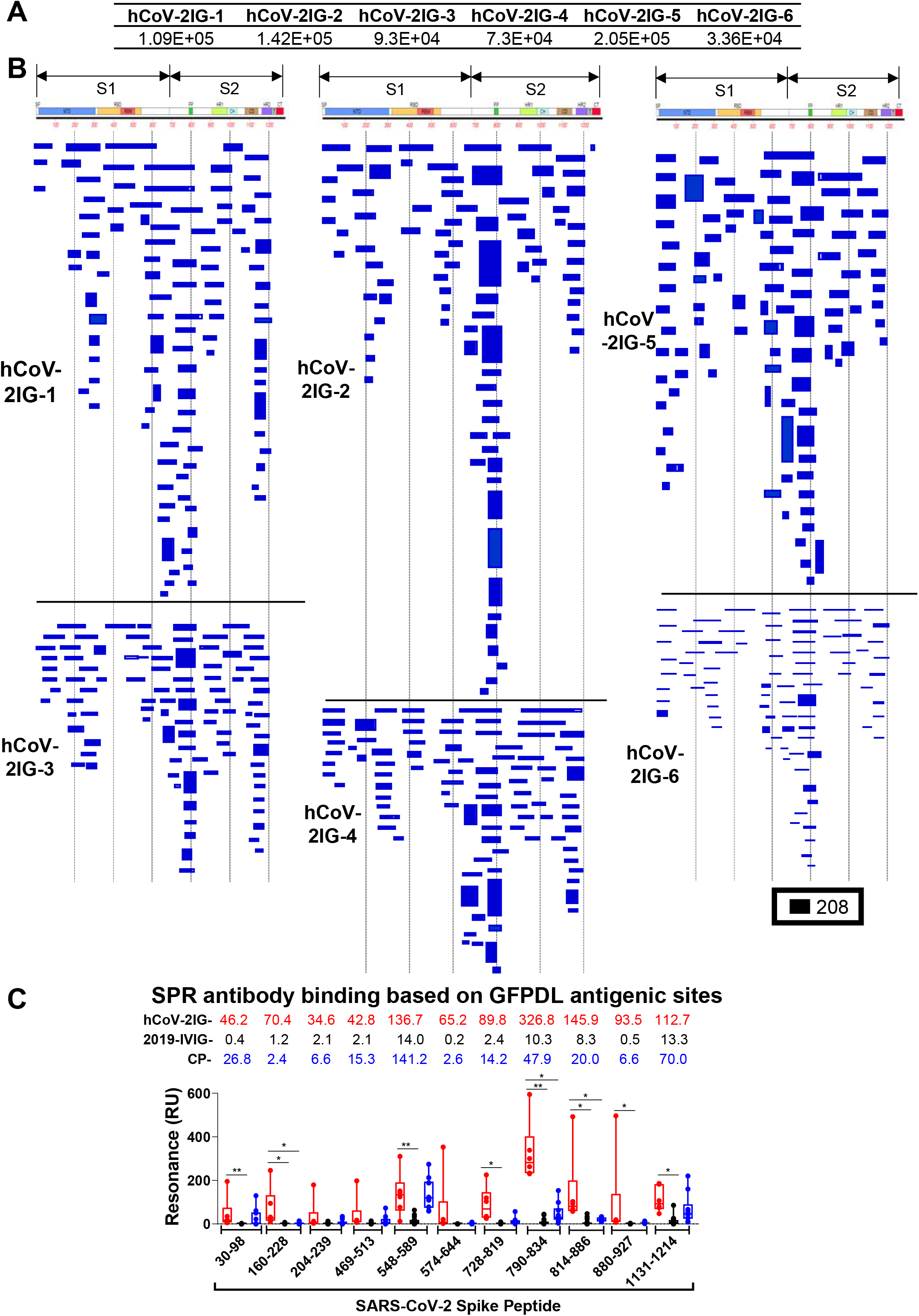
SARS-CoV-2 spike antibody epitope repertoires recognized by hCoV-2IG. SARS-CoV-2 spike GFPDL analyses of IgG antibodies in six batches of hCoV-2IG. (A) Number of IgG bound phage clones selected using SARS-CoV-2 spike GFPDL on six lots of hCoV-2IG (hCoV-2IG-1 to hCoV-2IG-6). (B) Epitope repertoires of IgG antibody in hCoV-2IG batches and their alignment to the spike protein of SARS-CoV-2. Graphical distribution of representative clones with a frequency of >2, obtained after affinity selection, are shown. The horizontal position and the length of the bars indicate the alignment of peptide sequence displayed on the selected phage clone to its homologous sequence in the SARS-CoV-2 spike. The thickness of each bar represents the frequency of repetitively isolated phage. Scale value is shown enclosed in a black box beneath the alignments. The GFPDL affinity selection data was performed in duplicate (two independent experiments by researcher in the lab, who was blinded to sample identity), and a similar number of phage clones and epitope repertoire was observed in both phage display analysis. (C) SPR binding of hCOV-2IG (n=6; in red), control pre-pandemic 2019-IVIG (n=16; in black) and convalescent plasma (n=9; in blue) with SARS-CoV-2 spike antigenic site peptides identified using GFPDL analysis in Fig. 1B. The amino acid designation is based on the SARS-CoV-2 spike protein sequence (Fig. S1). Total antibody binding is represented in maximum resonance units (RU) in this figure for 10-fold serum dilution of CP, and 1mg/mL of 2019-IVIG or hCoV-2IG. The numbers above the peptides show the mean value for each respective group antibody binding to the peptide and is color-coded (6 hCOV-2IG in red, 16 2019-IVIG in black, and 9 CPs in blue). All SPR experiments were performed twice and the researchers performing the assay were blinded to sample identity. The variations for duplicate runs of SPR was <4%. The data shown are average values of two experimental runs. The statistical significances between the hCoV-2IG vs 2019-IVIG vs CP for antibody binding to each peptide were performed using multiple group comparisons by non-parametric (Kruskal-Wallis) statistical test using Dunn’s post-hoc analysis in GraphPad prism. The differences were considered statistically significant with a 95% confidence interval when the p value was less than 0.05. (*, P values of ≤0.05, **, P values of ≤0.01).

### Neutralization capacity of CP and hCoV-2IG against the SARS-CoV-2 WA-1 and B.1.429, B1.1.7, P.1, B.1.351 VOCs

PsVNA was used to measure the neutralization activity of six hCoV-2 IG, nine CP, and 16 pre-pandemic 2019-IVIG lots against the predominant SARS-CoV-2 WA-1 strain and the VOCs currently spreading around the globe; U.S./CA (B.1.429), UK (B.1.1.7), JP (P.1), and SA (B.1.351). Both 50% (PsVNA50) and 80% (PsVNA80) neutralization titers were calculated. The spike proteins mutations in the VOCs used for production of the pseudovirions are shown in Table S2.

All sixteen pre-pandemic 2019-IVIG preparations demonstrated titers of <20 PsVNA50 against SARS-CoV-2 strains (Fig. 2A and Table S3). Among the nine CP lots tested against WA-1, variable PsVNA50 titers were observed, including one negative, one low (<1:80), six medium (>1:160<1:640) and two high (>1:640). In contrast, all six hCoV-2IG lots exhibited high PsVNA50 titers against WA-1 ranging between 1:1238-1:3309. PsVNA80 titers for hCoV-2IG ranged between 1:168-1:593, but none of the CP lots showed PsVNA80 titers above 1:80 (range <20 to 1:74) against WA-1 (Table S3). Neutralization of the VOCs showed gradual loss of titers as determined by either PsVNA50 or PsVNA80 for the hCoV-2IG and the CPs with greatest reduction in titers measured against the SA VOC (Fig. 2A and Table S3).

**Figure 2:**
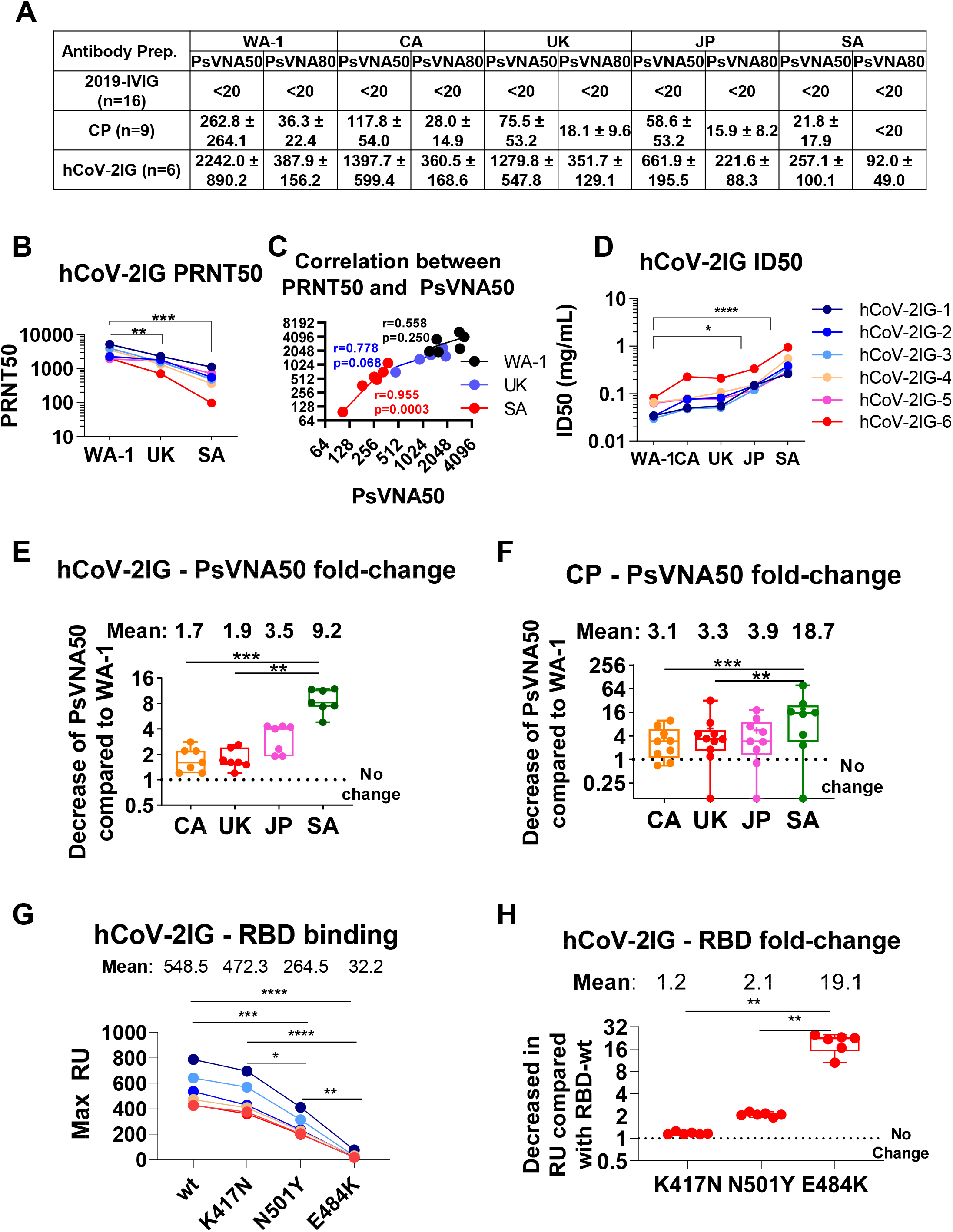
Neutralizing antibody titers and RBD binding antibodies of convalescent plasma and hCoV-2IG against various SARS-CoV-2 strains. (A) SARS-CoV-2 neutralizing antibody titers in CP, 2019-IVIG and hCoV-2IG preparations as determined by pseudovirus neutralization assay in 293-ACE2-TMPRSS2 cells with SARS-CoV-2 WA-1 strain, CA variant (B.1.429), UK variant (B.1.1.7), JP variant (P.1) or SA variant (B.1.351). PsVNA50 (50% neutralization titer) and PsVNA80 (80% neutralization titer) titers for control pre-pandemic 2019-IVIG (n=16), convalescent plasma (n =9) and hCoV-2IG (n = 6) were calculated with GraphPad prism version 8. Data show mean values + SEM for PsVNA50 and PsVNA80 titers for each of the 3 antibody groups against the SARS-CoV-2 WA-1, CA, UK, JP and SA variants. (B) End-point virus neutralization titers for six hCoV-2IG lots using wild type authentic SARS-CoV-2 WA-1, UK and SA virus strains in a classical BSL3 neutralization assay based on a plaque assay was performed as described in Materials and Methods. (C) Pearson two-tailed correlations are reported for the calculation of correlation of PRNT50 titers against wild-type SARS-CoV-2 strains (WA-1, UK or SA) and PsVNA50 titers against corresponding pseudovirions expressing either WA-1, UK or SA spike in pseudovirion neutralization assays for the six hCOV-2IG lots. (D) Antibody concentration (in mg/mL) required for each of the six hCoV-2IG batches to achieve 50% neutralization of SARS-CoV-2 WA-1, CA, UK, JP or SA variants in PsVNA. (E-F) Fold-decrease in PsVNA50 neutralization titers against emerging variant strain CA (B.1.429), UK (B.1.1.7), JP (P.1) and SA (B.1.351) for six hCoV-2IG lots (E) and nine CP lots (F) in comparison with SARS-CoV-2 WA-1 strain. The numbers above the group shows the mean fold-change for each variant. (G-H) Total antibody binding (Max RU) of 1mg/mL for the six batches of hCoV-2IG (hCoV-2IG-1 to hCoV-2IG-6) to purified WA-1 RBD (RBD-wt) and RBD mutants: RBD-K417N, RBD-N501Y and RBD-E484K by SPR (G). The numbers above the group show the mean antibody binding for each RBD. (H) Fold-decrease in antibody binding to mutants RBD-K417N, RBD-N501Y and RBD-E484K of hCoV-2IG in comparison with RBD-wt from WA-1 strain calculated from the data in Panel G. The numbers above the group shows the mean fold-change for each mutant RBD. All SPR experiments were performed twice and the researchers performing the assay were blinded to sample identity. The variations for duplicate runs of SPR was <5%. The data shown are average values of two experimental runs. The statistical significances between the variants for hCoV-2IG were performed using One-way ANOVA using Tukey’s pairwise multiple comparison test in GraphPad prism. The differences were considered statistically significant with a 95% confidence interval when the p value was less than 0.05. (*, P values of ≤0.05, **, P values of ≤0.01, ***, P values of ≤ 0.001, ****, P ≤0.0001).

For confirmation of the PsVNA neutralization titers, the six hCoV-2IG lots were also evaluated in a classical PRNT assay using VERO-E6 cells against authentic SARS-CoV-2 viruses representing WA-1 (USA-WA1/2020), UK-B1.1.7 (hCoV-19/England/204820464/2020), and SA-B1.351 (South Africa/KRISP-K005325/2020) strains (Fig. 2B and Table S4). Correlation of hCoV-2IG neutralization titers between PRNT50 and PsVNA50 were observed, as well as a similar decline in neutralization titers against the UK and SA VOC compared with WA-1 strain (Table S4 and Fig. 2B-C). Since all hCoV-2IG lots contain 100 mg/mL of IgG, it allowed calculation of ID50 values for the six hCoV-2IG lots (Table S5 and Fig 2D).

Compared to the WA-1 strain, the average PsVNA50 of the hCoV-2IG against CA, UK, JP and SA VOC were reduced by 1.7, 1.9, 3.5, and 9.2-fold respectively (Fig. 2E). Since the amount of SARS-CoV-2 specific IgG in CP lots is more variable and 5-10 fold lower compared with the hyperimmune hCoV-2IG, the CPs exhibited greater loss of neutralizing activities against the variants in comparison with hCoV-2IG. The average PsVNA50 of the CP against the CA, UK, JP, and SA VOC were reduced by 3.1, 3.3, 3.9, and 18.7-fold, respectively (Table S3 and Fig. 2F). PsVNA80 titers against the UK and JP VOC for all 9 CPs were lower (~2-fold) and were minimal or negligible against the SA VOC (Fig. 2A and Table S3). For hCoV-2IG, the PsVNA80 titers were similar for the WA-1, CA and UK strains, reduced by 1.75-fold against the JP variant, and decreased by 4.3-fold for SA VOC (Fig. 2A).

## Antibody binding of hCoV-2IG batches to RBD and RBD mutants: K417N, N501Y, and E484K

Many of the mutations in the spike protein of the different SARS-CoV-2 VOCs are unique (Fig. S2, Table S2), but a few key mutations among these strains are shared by VOCs as shown in Table S2. N501Y is shared among the UK, JP, and SA variants. E484K is shared between the JP and SA variants, and K417 is mutated to T in the JP variant, and to N in the SA variant. These key mutated residues have been shown to impact binding and neutralizing activity of antibodies in the post-infection and post-vaccination sera (*14*) (*15*). To further explore the possible contribution of these mutations in binding of hCoV-2IG batches, purified RBD proteins with individual mutations (K417N, N501Y, and E484K) were analyzed in SPR based antibody binding assays (Fig. 2G). The K417N had minimal to no impact on hCoV-2IG binding. The hCoV-2IG binding to RBD-N501Y was reduced by ~2-fold compared with WA-1 RBD. However, binding to RBD-E484K resulted in average 19-fold reduction in hCoV-2IG binding compared with the WA-1 RBD (Fig. 2H).

## DISCUSSION

In the current study we conducted in-depth analyses on six lots of hyperimmune globulin (hCoV-2IG) manufactured from plasma units collected from SARS-CoV-2 recovered individuals in 2020. Antibody epitope repertoire, neutralization of SARS-CoV-2 and VOCs, and binding to spike peptides as well as recombinant RBD expressing individual mutations observed in the VOC were evaluated in comparison with 9 convalescent plasma and 16 pre-pandemic 2019-IVIG.

The antibody epitope repertoires using SARS-CoV-2 spike GFPDL identified a diverse epitope fingerprint of both short and large antigenic sites spanning the entire spike protein. The hCoV-2IG antibodies most frequently bound to sites in the NTD, S1/S2 cleavage site, fusion peptide and heptad repeat domains in S2. In recent studies with monoclonal antibodies isolated from SARS-CoV-2 memory cells from COVID-19 patients, multiple neutralizing antibodies were identified that targeted the RBD, S1-NTD, S2, and S protein trimer(*16–18*). The most potent neutralizing antibodies that target directly the RBM/ACE2 interface were isolated at low frequency (*19*).

In light of the rapid spread of SARS-CoV-2 variants of concern around the globe, it is important to evaluate the therapeutic potential of both CP and hCoV-2IG against both early circulating SARS-CoV-2 strains and the emerging VOCs (*20–22*) that can define the therapeutic potential of these antibody preparations. In the current study, all six hCoV-2IG lots demonstrated a small decline in neutralization titers (and increase in ID50) against CA, UK (~ 2-fold) followed by JP (~3.5-fold) VOC, and a significant decrease in neutralization activity against SA VOC (~9-fold). The nine CP evaluated demonstrated a range of neutralization titers compared with the hCoV-2IG against the WA-1 strain, and a larger reduction in PsVNA50 titers against the VOCs compared with hCoV-2IG.

Most of the SARS-CoV-2 VOCs that have been spreading in different parts of the world have multiple mutations both in the spike and other genes. However, several VOCs share one or more mutations in the RBD. Decrease in antibody binding to the RBD interface with ACE2 receptor is probably the key reason for loss of SARS-CoV-2 neutralizing activity against the VOCs. Interestingly, while most of the interactions between the Receptor Binding Motif (RBM) 25 residues and the predominant IGHV3-53/IGHV3-66 neutralizing antibodies are mediated by hydrogen bonds. Only K417 and E484 have been described to form a salt bridge resulting in a stronger interaction and higher immune pressure (*23*). We found that the E484K mutation, which is shared between Brazil/JP P.1 VOC and SA B.1.351 VOC significantly reduced binding of hCoV-2IG to the RBD (19-fold reduction) compared with RBD-wt. In contrast, the K417N had only minimal effect on RBD binding and the N501Y reduced binding of the hCoV-2IG by 2-fold. Therefore, virus neutralization may be impacted both by specific amino acid mutations in the RBD/RBM and by the specificity of the polyclonal antibodies that bind to other sites on the SARS-CoV-2 spike.

The correlate of protection in terms of antibody neutralizing titers has not been identified in ongoing vaccine trials. However, studies in rhesus macaques showed that passive transfer of 250 mg/kg SARS-CoV-2 IgG one day after challenge, reduced the peak lung viral loads and cleared the virus by day 3 (*24*). Convalescent plasma demonstrated significant loss of neutralizing activities against the emerging VOC, especially the SA B.1.351(*14*). However, some reduction in post-vaccination titers was observed against the new variants, especially the South African VOC, but SARS-CoV-2 vaccines that elicit high and durable neutralization titers may still be effective against severe disease associated with VOCs (*15*).

Our study underscores the advantage of using hyperimmune immunoglobulin products (hCoV-2IG) compared with CP for treatment of SARS-CoV-2 infected patients. The neutralizing titers were found to decline significantly after 3 or 4 months in recovered COVD-19 patients (*25*). Therefore, screening of multiple CP units prior to pooling for fractionation can ensure high neutralizing titer hCoV-2 IG products and lot to lot consistency. The added values of hCoV-2IG over CP is even more critical in the face of emerging more transmissible VOCs that are spreading in several countries around the globe. Casadevall et al. emphasized that antibody preparations should contain sufficiently high concentrations of specific immunoglobulin to mediate biological effect against SARS-CoV-2 and its variants and should be administered early post-exposure (*26*).

In summary, both CP and hCoV-2IG demonstrated reduced neutralization titers ranging from ~2-4 fold against UK & JP VOCs and ~10-20 fold against SA VOC. Our findings indicate that treatment of COVID-19 patients with hCoV-2IG/CP may still be feasible but would likely require administration of higher volumes, or repeated infusions, in patients infected with the emerging SARS-CoV-2 variants. This will require rapid development of RT-PCR based diagnostics or other diagnostic assays that are designed to differentiate the B.1.1.7 and B.1.351 variants, and other emerging SARS-CoV-2 variants like the California (B.1.429) and Japan (P.1) strains. Furthermore, in countries where a new VOC becomes dominant, the manufacturing of new hCoV-2IG should incorporate screening of the plasma and of the hCoV-2IG lots for neutralization activities against VOC. This study suggests that in countries with multiple co-circulating SARS-CoV-2 variants, the identification of the infecting SARS-CoV-2 strain prior to treatment with hCoV-2IG will be critical in determining the effectiveness of antibody therapy in COVID-19 patients.

## METHODS

### Study design

The objective of this study was to investigate antibody binding and neutralizing capacity of various therapeutic polyclonal CP or purified hCoV-2-IG antibody preparations being evaluated in the clinical trials with important emerging SARS-CoV-2 variant of concern (VOC). Such variants of concern (VOC) are the United Kingdom (UK) variant B.1.1.7, California (CA) variant B.1.429, Japan (JP) variant P.1, and the South Africa (SA) variant B.1.351. There are multiple IVIG products approved by the FDA. These are polyclonal antibodies made from U.S. plasma donors. Each lot of product is derived from 10,000 or more donors. The manufacturing processes vary between manufacturers and usually include cold alcohol fractionation (Cohn-Oncley), anion-exchange and size-exclusion chromatography. Sixteen intravenous immunoglobulin (IVIG) batches were produced from plasma collected in 2019 (each lot derived from >10,000 donors) from five manufacturers, prior to August 2019. The final product is sterile-filtered IgG (> 95%) and formulated at 100 mg/mL Nine random CP lots were obtained from recovered COVID-19 patients. Six hCoV-2IG batches prepared from 250-400 COVID-19 CP donors per lot were obtained from three commercial companies. This study was approved by the Food and Drug Administration’s Research Involving Human Subjects Committee (RIHSC #2020-04-02).

### Lentivirus pseudovirion neutralization assay (PsVNA)

Antibody preparations were evaluated by SARS-CoV-2 pseudovirus neutralization assay (PsVNA) using WA-1 strain, UK variant (B.1.1.7 with spike mutations: H69-V70del, Y144del, N501Y, A570D, D614G, P681H, T716I, S982A, and D1118H), SA variant (B.1.351 strain with spike mutations L18F, D80A, D215G, L242-244del, R246I, K417N, E484K, N501Y, D614G, and A701V), CA variant (B.1.429 strain with spike mutations S13I, W152C, L452R, D614G) and JP variant (P.1 strain with spike mutations L18F, T20N, P26S, D138Y, R190S, K417T, E484K, N501Y, H655Y, T1027I, D614G, V1176F) (Table S1). The PsVNA using 293-ACE2-TMPRSS2 cell line was described previously (*10, 11*).

Briefly, human codon-optimized cDNA encoding SARS-CoV-2 S glycoprotein of the WA-1, UK VOC, CA VOC, JP VOC and SA VOC was synthesized by GenScript and cloned into eukaryotic cell expression vector pcDNA 3.1 between the *BamHI* and *XhoI* sites. Pseudovirions were produced by co-transfection Lenti-X 293T cells with psPAX2(gag/pol), pTrip-luc lentiviral vector and pcDNA 3.1 SARS-CoV-2-spike-deltaC19, using Lipofectamine 3000. The supernatants were harvested at 48h post transfection and filtered through 0.45μm membranes and titrated using 293T-ACE2-TMPRSS2 cells (HEK293T cells that express ACE2 and TMPRSS2 proteins).

For the neutralization assay, 50 μL of SARS-CoV-2 S pseudovirions were pre-incubated with an equal volume of medium containing serum at varying dilutions at room temperature for 1 h, then virus-antibody mixtures were added to 293T-ACE2-TMPRSS2 cells in a 96-well plate. The input virus with all three SARS-CoV-2 strains used in the current study were the same (2x 10^5^ Relative light units/50 μL/well). After a 3 h incubation, the inoculum was replaced with fresh medium. Cells were lysed 24 h later, and luciferase activity was measured using luciferin. Controls included cells only, virus without any antibody and positive sera. The cut-off value or the limit of detection for neutralization assay is 1:10.

### Classical wild-type SARS-CoV-2 virus neutralization assay

100 TCID50 of SARS-CoV-2 WA-1 (USA-WA1/2020), UK-B1.1.7 (hCoV-19/England/204820464/2020), and SA-B1.351 (South Africa/KRISP-K005325/2020) strains was incubated with 2-fold serial dilutions in a round bottom plate at 37°C for 1 hr. The virus-antibody mixture was then added to a 96-well plate with 5×10^4^ Vero E6 cells. After 1 h the mixture was removed and replenished with fresh MEM containing 2% FBS. Cells were incubated at 37°C for an additional 48 hours, then fixed with 4% paraformaldehyde, followed by staining of cells with 0.1% crystal violet in 20% methanol. The PRNT50 and PRNT90 titers were calculated as the last serum dilution resulting in at least 50% and 90% SARS-CoV-2 neutralization, respectively.

### Proteins

Recombinant SARS-CoV-2 spike receptor binding domain (RBD) and its mutants were purchased from Sino Biologicals (RBD-wt; 40592-V08H82, RBD-K417N; 40592-V08H59, RBD-N501Y; 40592-V08H82 and RBD-E484K; 40592-V08H84). Recombinant purified RBD proteins used in the study were produced in 293 mammalian cells. The native receptor-binding activity of the spike RBD proteins was determined by binding to the 5 μg/mL human ACE2 protein(*10–12*).

### SARS-CoV-2 Gene Fragment Phage Display Library (GFPDL) construction

DNA encoding the spike gene of SARS-CoV-2 isolate Wuhan-Hu-1 strain (GenBank: MN908947.3) was chemically synthesized and used for cloning. A gIII display-based phage vector, fSK-9-3, was used where the desired polypeptide can be displayed on the surface of the phage as a gIII-fusion protein. Purified DNA containing spike gene was digested with *DNaseI* to obtain gene fragments of 50-1500 bp size range (18 to 500 amino acids) and used for GFPDL construction as described previously (*10–12*).

### Affinity selection of SARS-CoV-2 GFPDL phages

Prior to panning of GFPDL with polyclonal hCoV-2IG antibodies, Ig components, which could non-specifically interact with phage proteins, were removed by incubation with UV-killed M13K07 phage-coated Petri dishes. Equal volumes of each of the six hCoV-2IG lots were used for GFPDL panning. GFPDL affinity selection was carried out in-solution with protein A/G resin as previously described(*10–12*). Briefly, the hCoV-2IG lot was incubated with the GFPDL and the protein A/G resin, the unbound phages were removed by PBST (PBS containing 0.1 % Tween-20) wash followed by PBS. Bound phages were eluted by addition of 0.1 N Gly-HCl pH 2.2 and neutralized by adding 8 μL of 2 M Tris solution per 100 μL eluate. After panning, antibody-bound phage clones were amplified, the inserts were sequenced, and the sequences were aligned to the SARS-CoV-2 spike gene, to define the fine epitope specificity in these polyclonal hCoV-2IG lots.

The GFPDL affinity selection was performed in duplicate (two independent experiments by research fellow in the lab, who was blinded to sample identity). Similar numbers of bound phage clones and epitope repertoire were observed in the two GFPDL panning.

### Antibody binding kinetics to SARS-CoV-2 RBD mutants or SARS-CoV-2 peptides by Surface Plasmon Resonance (SPR)

Steady-state equilibrium binding of hCoV-2IG lots was monitored at 25°C using a ProteOn surface plasmon resonance (BioRad). The purified recombinant SARS-CoV-2 RBD proteins were captured to a Ni-NTA sensor chip with 200 resonance units (RU) in the test flow channels. The protein density on the chip was optimized such as to measure monovalent interactions independent of the antibody isotype (*10–12, 27*). The biotinylated SARS-CoV-2 peptides were captured on NLC chip and used for peptide antibody profiling of hCoV-2IG, CP and 2019-IVIG lots.

Serial dilutions (1 mg/mL, 0.33 mg/mL and 0.11 mg/mL) of freshly prepared hCoV-2IG or 2019-IVIG or 10-fold dilution of CP in BSA-PBST buffer (PBS pH 7.4 buffer with Tween-20 and BSA) were injected at a flow rate of 50 μL/min (120 sec contact duration) for association, and disassociation was performed over a 600-second interval. Responses from the protein surface were corrected for the response from a mock surface and for responses from a buffer-only injection. SPR was performed with serially diluted samples in this study. Total antibody binding was calculated with BioRad ProteOn manager software (version 3.1). All SPR experiments were performed twice, and the researchers performing the assay were blinded to sample identity. The maximum resonance units (Max RU) data shown in the figures were the calculated RU signal for the 1 mg/mL hCoV-2IG sample or 2019-IVIG or 10-fold dilution of CP.

### Statistical Analysis

All experimental data were analyzed in GraphPad Prism, version 9.0.1 (GraphPad software Inc, San Diego, CA) or R package. Differences between groups were analyzed using multiple group comparisons by non-parametric (Kruskal-Wallis) statistical test using Dunn’s post-hoc analysis. The difference within each group were performed using one-way ANOVA using Tukey’s pairwise multiple comparison test. The differences were considered statistically significant with a 95% confidence interval when the p value was less than 0.05. (*, P values of ≤0.05, **, P values of ≤0.01, ***, P values of ≤ 0.001, ****, P ≤0.0001). Correlation analysis of PRNT and PsVNA titers were performed by computing Pearson’s correlation coefficient in Graphpad.

## Ethics Statement

This study was approved by Food and Drug Administration’s Research Involving Human Subjects Committee (RIHSC #2020-04-02). All assays performed fell within the permissible usages in the original consent.

## ACKNOWLEDGEMENTS

We thank Keith Peden and Marina Zaitseva for their insightful review of the manuscript. We thank Carol Weiss for providing plasmid clones expressing SARS-CoV-2 variants and Dorothy Scott for providing convalescent plasma and hCoV-2IG.

## Role of Funder

The research work described in this manuscript was supported by FDA intramural funds and NIH-NIAID IAA #AAI20040. The funders had no role in study design, data collection and analysis, decision to publish, or preparation of the manuscript.

The content of this publication does not necessarily reflect the views or policies of the Department of Health and Human Services, nor does mention of trade names, commercial products, or organizations imply endorsement by the U.S. Government.

## AUTHOR CONTRIBUTIONS

**Designed research:** S.K.

**Clinical specimens and clinical data:** H.G. and B.G.

**Performed research:** J.T., Y.L., S.R., G.G., C. H., S.C., T.W., and S.K.

**Contributed to Writing:** S.K., H.G. and B.G.

**Conflict of Interest Disclosures:** The authors declare no competing interests.

## Data and Materials availability

All data needed to evaluate the conclusions in the current study are present in the main figures and/or the Supplementary Materials. The materials generated during the current study are available from corresponding author under a material transfer agreement on reasonable request.

## SUPPLEMENTAL INFORMATION

Supplementary Figure 1: SARS-CoV-2 epitope profile of six hCoV-2IG lots in pseudovirion neutralization assay.

Supplementary Figure 2. Multiple sequence alignment of Spike protein of SARS-CoV-2 variants.

Supplementary Table 1: Sequence conservation of GFPDL-identified antigenic regions/sites among different SARS-CoV-2 VOCs.

Supplementary Table 2: SARS-CoV-2 variants mutations introduced in the spike plasmid for production of SARS-CoV-2 pseudovirions to test them in PsVNA.

Supplementary Table 3: Neutralization titers of convalescent plasma, IVIG and hCoV-2IG against SARS-CoV-2 variants.

Supplementary Table 4: PRNT50 and PRNT80 of the six hCoV-2IG batches against SARS-CoV-2 WA-1 and UK & SA VOCs in classical wild-type SARS-CoV-2 virus neutralization assay.

Supplementary Table 5: Antibody concentration (in mg/mL) required for each of the six hCoV-2IG batches to achieve 50% (ID50) or 80% (ID80) neutralization of SARS-CoV-2 variants in PsVNA.

---

**Supplementary Figure 1.**
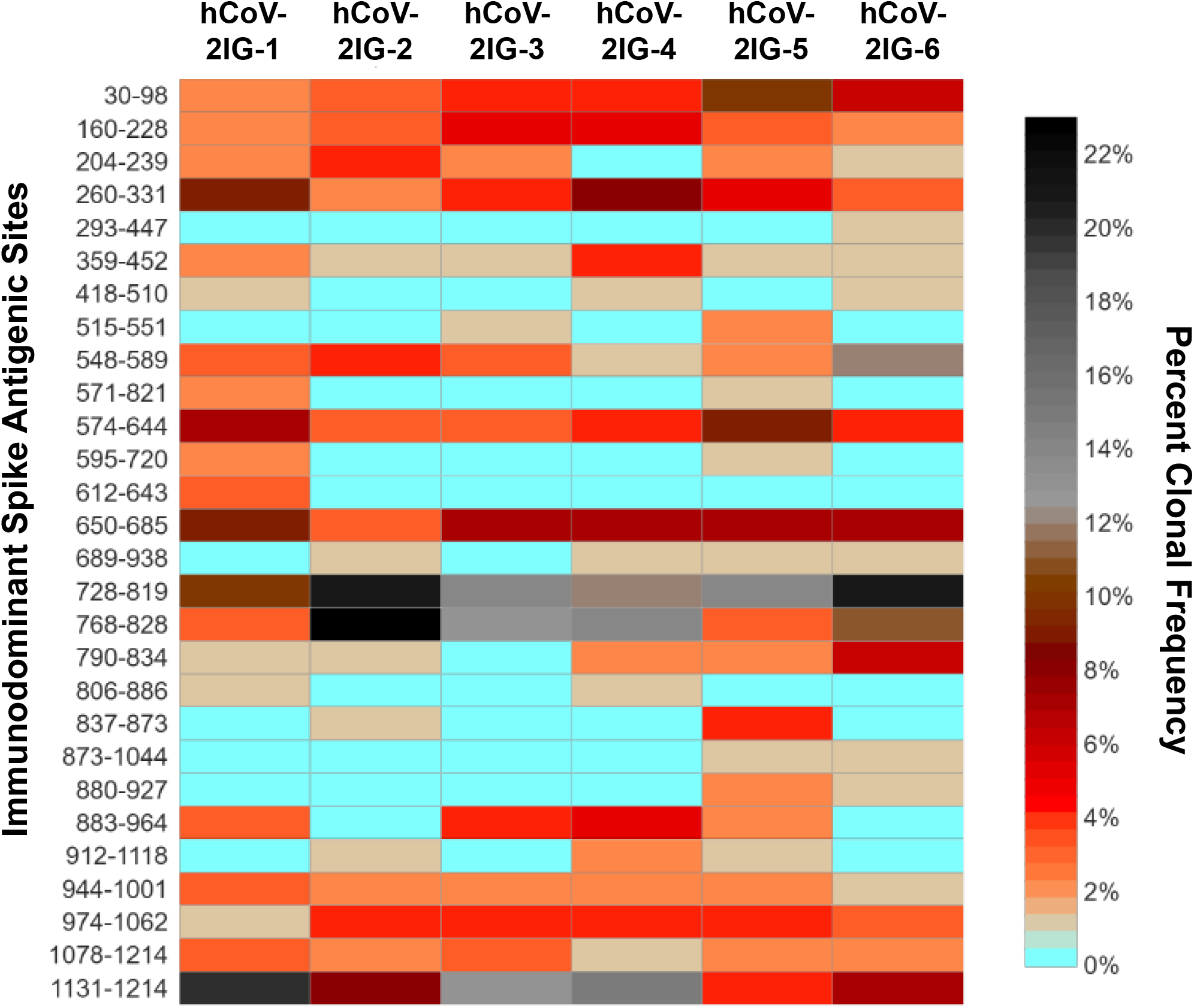
SARS-CoV-2 epitope profile of six hCoV-2IG batches. Heat map of immunodominant sites (≥3 clonal frequency in at least one hCoV-2IG lot) on the SARS-CoV-2 spike recognized by IgG antibodies in six hCoV-2IG lots identified using GFPDL analyses. The immunodominant sites on the left indicate amino acid residue of the antigenic sites in the spike protein. Color scale on the right represents range of percentage of clonal occurrences (frequency) of each site. Heat map was generated using R package.

**Supplementary Figure S2.**
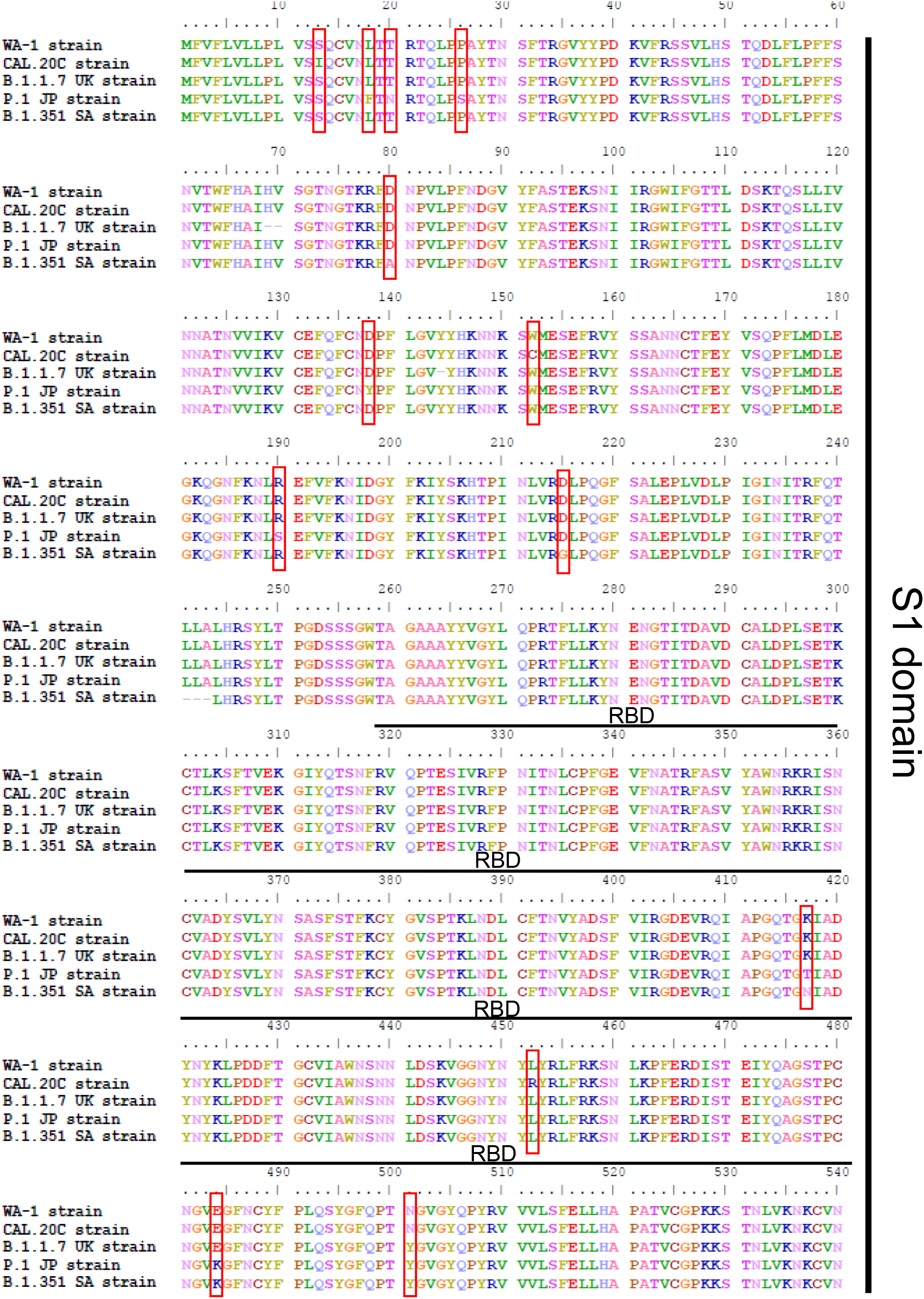
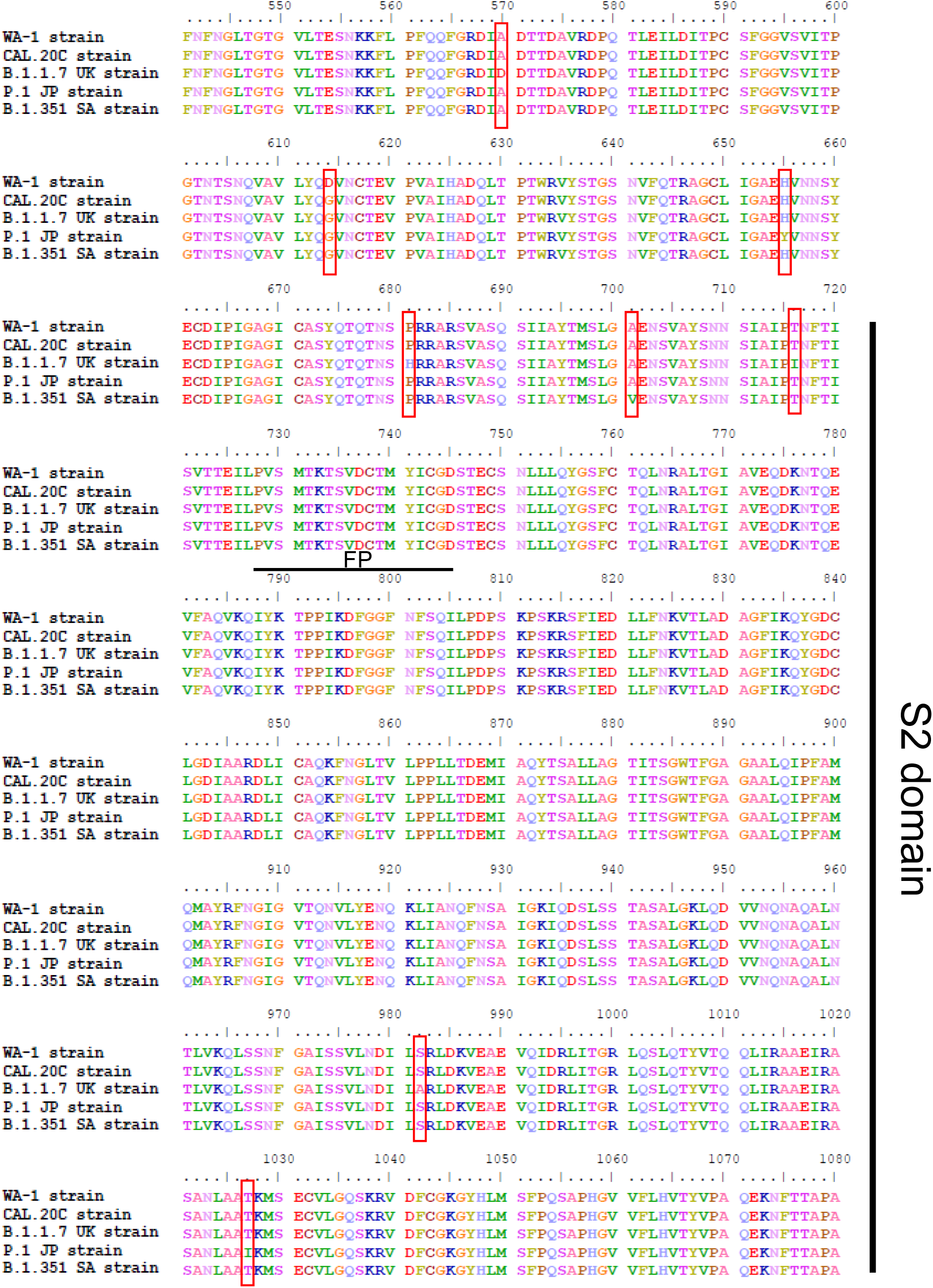
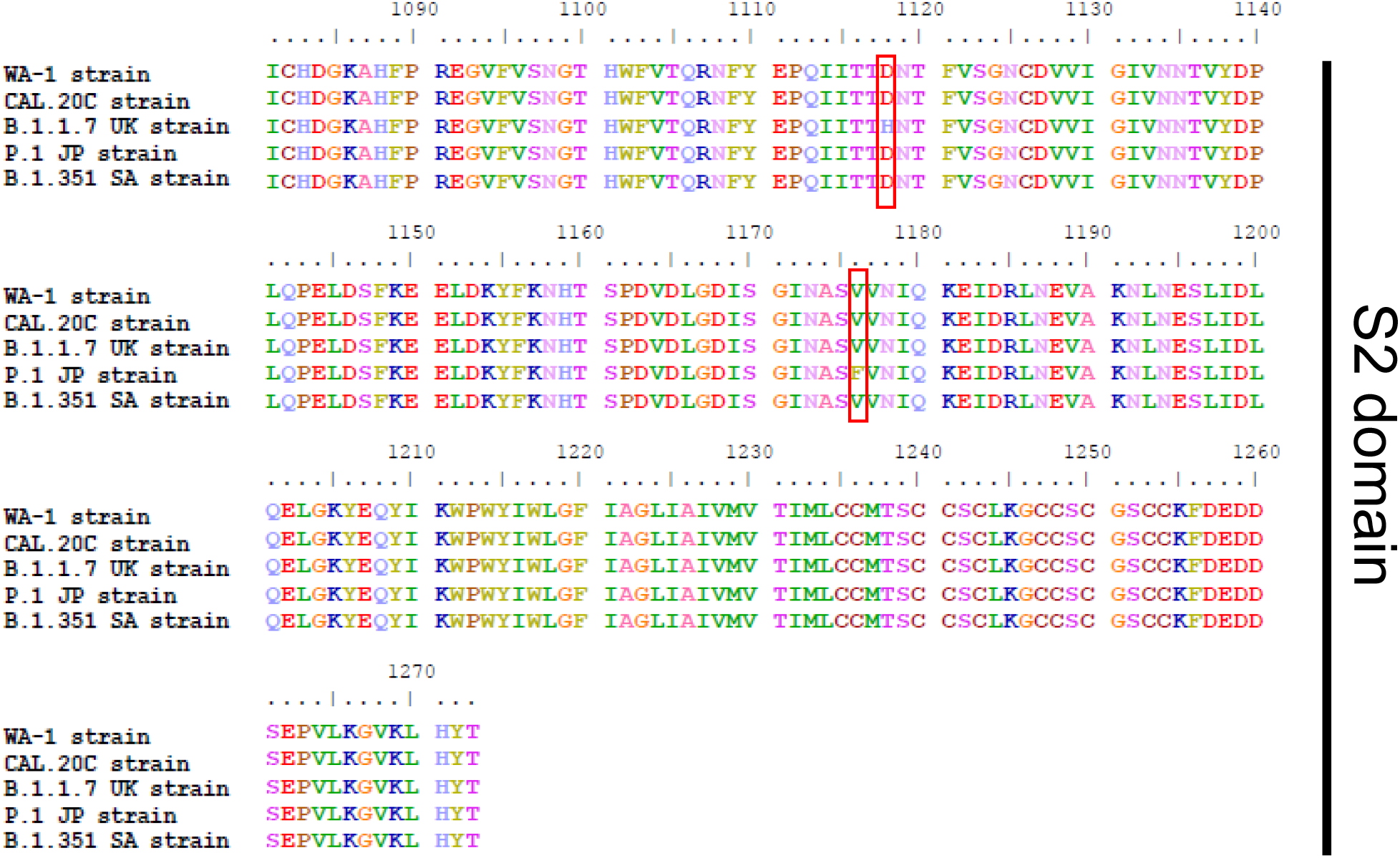
Multiple sequence alignment of Spike protein of SARS-CoV-2 variants. Multiple sequence alignment of various SARS-CoV-2 variants namely WA-1 strain (QII87782.1), CA variant (B.1.429, EPI_ISL_648527), UK variant (B.1.1.7, QQQ47833.1), JP variant (P.1, QRX39425.1), and SA variant (B.1.351, EPI_ISL_678597) was performed using MAFFT version 7 alignment tool (https://mafft.cbrc.jp/alignment/software/). Mutations in any or all of the variants are indicated with a red outline around each of them. Various domains of the spike protein are also indicated namely S1, S2, RBD and FP domains.

**Supplementary Table 1:**
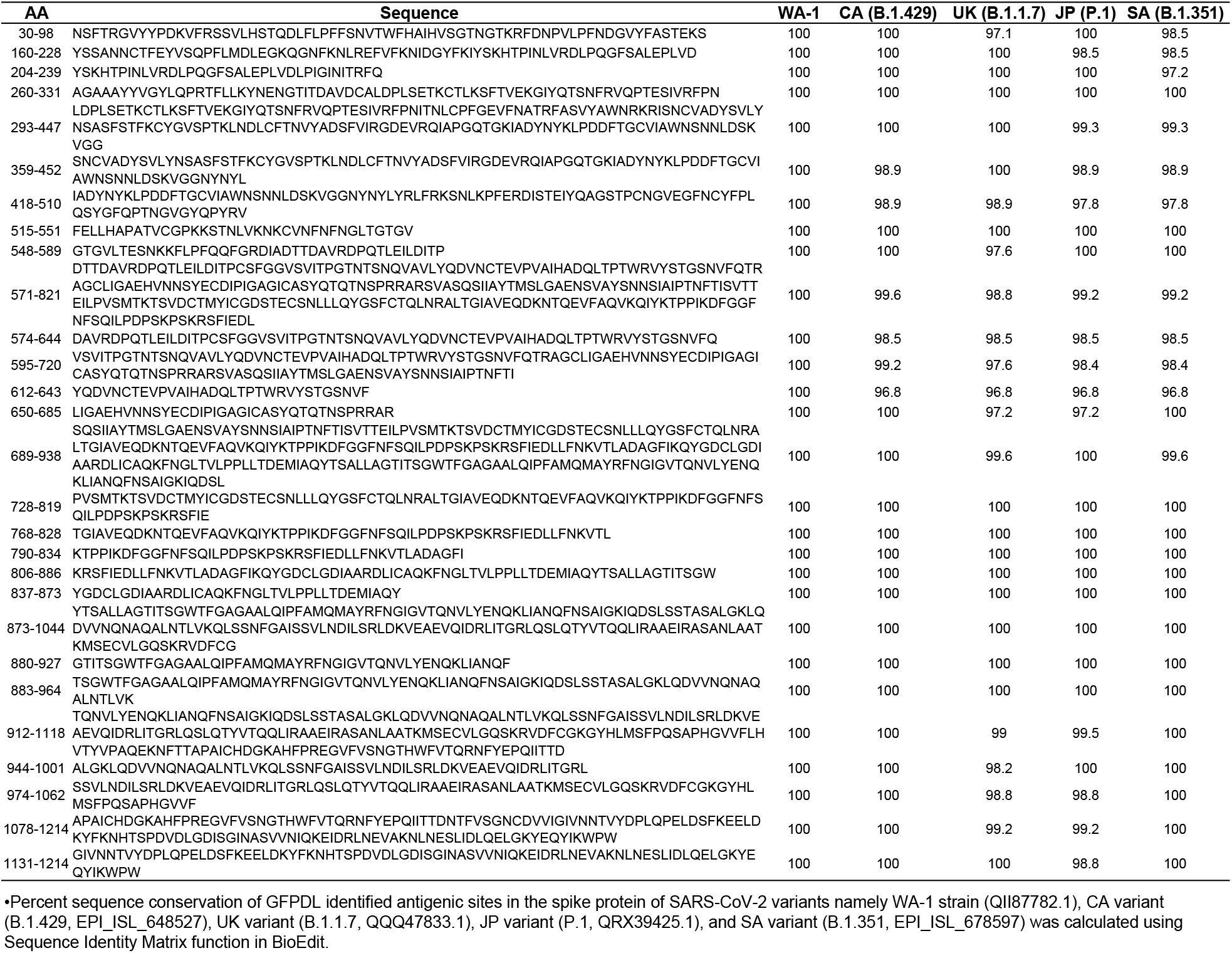
Sequence conservation of GFPDL-identified antigenic regions/sites among different SARS-CoV-2 VOCs*

**Supplementary Table 2:** SARS-CoV-2 variants mutations introduced in the spike plasmid for production of SARS-CoV-2 pseudovirions to test them in PsVNA.

| **Mutations constructed in the spike plasmids** |  |
| --- | --- |
| **CA** | S13I, W152C, L452R, D614G |
| **UK** | Del69-70, Del145,N501Y, A570D, D614G, P681H, T716I, S982A, D1118H |
| **JP** | L18F, T20N, P26S, D138Y, R190S, K417T, E484K, N501Y, H655Y, T1027I, D614G, V1176F |
| **SA** | L18F, D80A, D215G, Del242-244, K417N, E484K, N501Y, D614G, A701V |

**Supplementary Table 3:** Neutralization titers of convalescent plasma, I2019-VIG and hCoV-2IG against SARS-CoV-2 variants*

|  | WA-1 |  | CA |  | UK |  | JP |  | SA |  |
| --- | --- | --- | --- | --- | --- | --- | --- | --- | --- | --- |
|  | PsVNA50 | PsVNA80 | PsVNA50 | PsVNA80 | PsVNA50 | PsVNA80 | PsVNA50 | PsVNA80 | PsVNA50 | PsVNA80 |
| **IVIG batches produced in 2019 prior to COVID-19** |  |  |  |  |  |  |  |  |  |  |
| 2019-IVIG-1 | <20 | <20 | <20 | <20 | <20 | <20 | <20 | <20 | <20 | <20 |
| 2019-IVIG-2 | <20 | <20 | <20 | <20 | <20 | <20 | <20 | <20 | <20 | <20 |
| 2019-IVIG-3 | <20 | <20 | <20 | <20 | <20 | <20 | <20 | <20 | <20 | <20 |
| 2019-IVIG-4 | <20 | <20 | <20 | <20 | <20 | <20 | <20 | <20 | <20 | <20 |
| 2019-IVIG-5 | <20 | <20 | <20 | <20 | <20 | <20 | <20 | <20 | <20 | <20 |
| 2019-IVIG-6 | <20 | <20 | <20 | <20 | <20 | <20 | <20 | <20 | <20 | <20 |
| 2019-IVIG-7 | <20 | <20 | <20 | <20 | <20 | <20 | <20 | <20 | <20 | <20 |
| 2019-IVIG-8 | <20 | <20 | <20 | <20 | <20 | <20 | <20 | <20 | <20 | <20 |
| 2019-IVIG-9 | <20 | <20 | <20 | <20 | <20 | <20 | <20 | <20 | <20 | <20 |
| 2019-IVIG-10 | <20 | <20 | <20 | <20 | <20 | <20 | <20 | <20 | <20 | <20 |
| 2019-IVIG-11 | <20 | <20 | <20 | <20 | <20 | <20 | <20 | <20 | <20 | <20 |
| 2019-IVIG-12 | <20 | <20 | <20 | <20 | <20 | <20 | <20 | <20 | <20 | <20 |
| 2019-IVIG-13 | <20 | <20 | <20 | <20 | <20 | <20 | <20 | <20 | <20 | <20 |
| 2019-IVIG-14 | <20 | <20 | <20 | <20 | <20 | <20 | <20 | <20 | <20 | <20 |
| 2019-IVIG-15 | <20 | <20 | <20 | <20 | <20 | <20 | <20 | <20 | <20 | <20 |
| 2019-IVIG-16 | <20 | <20 | <20 | <20 | <20 | <20 | <20 | <20 | <20 | <20 |
| **Convalescent plasma batches produced from COVID-19 survivors** |  |  |  |  |  |  |  |  |  |  |
| CP-1 | 243.1 | 28.0 | 24.7 | <20 | 53.4 | <20 | 287.0 | 61.1 | 107.4 | 30.6 |
| CP-2 | 369.2 | 56.8 | 122.9 | 30.0 | 128.3 | 23.9 | 126.0 | 24.7 | 25.5 | <20 |
| CP-3 | 204.9 | 30.1 | 146.9 | 37.3 | 168.4 | 24.0 | 112.0 | 27.3 | 47.2 | <20 |
| CP-4 | 268.0 | 31.3 | 92.8 | <20 | 59.8 | <20 | 53.8 | <20 | <20 | <20 |
| CP-5 | <20 | <20 | <20 | <20 | <20 | <20 | <20 | <20 | <20 | <20 |
| CP-6 | 211.1 | 47.2 | 123.3 | 26.1 | 64.8 | 21.6 | <20 | <20 | <20 | <20 |
| CP-7 | 67.0 | <20 | 98.5 | 22.2 | 36.6 | <20 | 24.1 | <20 | <20 | <20 |
| CP-8 | 119.6 | 30.2 | 156.8 | 33.0 | 35.4 | <20 | <20 | <20 | <20 | <20 |
| CP-9 | 852.7 | 74.9 | 191.0 | 55.6 | 100.7 | 35.4 | 122.6 | 25.4 | 51.7 | <20 |
| **hCoV-2IG batches produced from COVID-19 convalescent plasma donors** |  |  |  |  |  |  |  |  |  |  |
| hCoV-2IG-1 | 2878.8 | 515.0 | 2009.3 | 613.7 | 1799.4 | 401.1 | 672.2 | 269.1 | 382.0 | 154.5 |
| hCoV-2IG-2 | 2919.7 | 435.0 | 1301.5 | 305.8 | 1224.9 | 394.0 | 681.7 | 235.8 | 259.0 | 119.5 |
| hCoV-2IG-3 | 3309.1 | 593.2 | 2081.3 | 495.7 | 1957.1 | 499.1 | 833.4 | 302.2 | 278.0 | 124.1 |
| hCoV-2IG-4 | 1514.5 | 292.9 | 1257.8 | 261.8 | 935.6 | 286.8 | 653.3 | 176.7 | 184.5 | 80.5 |
| hCoV-2IG-5 | 1590.1 | 323.0 | 1295.7 | 341.2 | 1288.5 | 402.4 | 832.3 | 280.8 | 333.6 | 39.7 |
| hCoV-2IG-6 | 1239.5 | 168.3 | 440.8 | 144.4 | 473.4 | 126.8 | 298.5 | 65.2 | 105.6 | 33.5 |

\*PsVNA titer Cut-off value: 1:20.

**Supplementary Table 4:** PRNT50 and PRNT80 of the six hCoV-2IG batches against SARS-CoV-2 WA-1, UK and SA strains in classical wild-type SARS-CoV-2 virus neutralization assay.

|  | WA-1 |  | UK |  | SA |  |
| --- | --- | --- | --- | --- | --- | --- |
|  | PRNT50 | PRNT90 | PRNT50 | PRNT90 | PRNT50 | PRNT90 |
| **hCoV-2IG-1** | 5184.7 | 1723.8 | 2324.7 | 730.1 | 1111.4 | 296.1 |
| **hCoV-2IG-2** | 2260.6 | 264.4 | 1826.4 | 405.6 | 553.7 | 200.3 |
| **hCoV-2IG-3** | 4030 | 1504 | 1556.8 | 463.5 | 486.4 | 130.5 |
| **hCoV-2IG-4** | 3580.7 | 1153.8 | 1336.9 | 298.6 | 361 | 79.3 |
| **hCoV-2IG-5** | 1929.1 | 803.8 | 1674.6 | 370.8 | 716 | 194.3 |
| **hCoV-2IG-6** | 1988.8 | 663.7 | 708.1 | 50.9 | 96.8 | 4.3 |

**Supplementary Table 5:** Antibody concentration (in mg/mL) required for each of the six hCoV-2IG batches to achieve 50% (ID50) or 80% (ID80) neutralization of SARS-CoV-2 variants in PsVNA.

|  | WA-1 |  | CA |  | UK |  | JP |  | SA |  |
| --- | --- | --- | --- | --- | --- | --- | --- | --- | --- | --- |
|  | ID50<br>(mg/mL) | ID80<br>(mg/mL) | ID50<br>(mg/mL) | ID80<br>(mg/mL) | ID50<br>(mg/mL) | ID80<br>(mg/mL) | ID50<br>(mg/mL) | ID80<br>(mg/mL) | ID50<br>(mg/mL) | ID80<br>(mg/mL) |
| hCoV-2IG-1 | 0.035 | 0.194 | 0.050 | 0.163 | 0.056 | 0.249 | 0.149 | 0.372 | 0.262 | 0.647 |
| hCoV-2IG-2 | 0.034 | 0.230 | 0.077 | 0.327 | 0.082 | 0.254 | 0.147 | 0.424 | 0.386 | 0.837 |
| hCoV-2IG-3 | 0.030 | 0.169 | 0.048 | 0.202 | 0.051 | 0.200 | 0.120 | 0.331 | 0.360 | 0.806 |
| hCoV-2IG-4 | 0.066 | 0.341 | 0.080 | 0.382 | 0.107 | 0.349 | 0.153 | 0.566 | 0.542 | 1.242 |
| hCoV-2IG-5 | 0.063 | 0.310 | 0.077 | 0.293 | 0.078 | 0.249 | 0.120 | 0.356 | 0.300 | 2.519 |
| hCoV-2IG-6 | 0.081 | 0.594 | 0.227 | 0.692 | 0.211 | 0.789 | 0.335 | 1.534 | 0.947 | 2.988 |

## Notes

### Competing Interest Statement

The authors have declared no competing interest.

